# Proteasome-mediated regulation of Cdhr1a by Siah1 modulates photoreceptor development and survival in zebrafish

**DOI:** 10.1101/2020.05.15.098350

**Authors:** Warlen P. Piedade, Kayla Titialii-Torres, Ann Morris, Jakub Famulski

**Affiliations:** Department of Biology, University of Kentucky

**Keywords:** CDHR1a, Siah E3 ubiquitin ligase, photoreceptor, zebrafish, UPS, cell death, retina

## Abstract

Congenital retinal dystrophies are a major cause of unpreventable and incurable blindness worldwide. Mutations in CDHR1, a retina specific cadherin, are associated with cone-rod dystrophy. The ubiquitin proteasome system (UPS) is responsible for mediating orderly and precise targeting of protein degradation to maintain biological homeostasis and coordinate proper development, including retinal development. Recently, our lab uncovered that the seven in absentia (Siah) family of E3 ubiquitin ligases play a role in optic fissure fusion, and identified Cdhr1a as a potential target of Siah. Using two-color whole mount *in situ* hybridization and immunohistochemistry, we detected *siah1* and *cdhr1a* co-expression as well as protein co-localization in the retinal outer nuclear layer (ONL), and more precisely in the connecting cilium of rods and cones between 3-5 days post fertilization (dpf). We confirmed that Siah1 targets Cdhr1a for proteasomal degradation by co-transfection and co-immunoprecipitation in cell culture. To analyze the functional importance of this interaction, we created two transgenic zebrafish lines that express *siah1* or an inactive *siah1* (*siah1*ΔRING) under the control of the heat shock promoter to modulate Siah activity during photoreceptor development. Overexpression of *siah1*, but not *siah1*ΔRING, resulted in a decrease in the number of rods and cones at 72 hours post fertilization (hpf). The number of retinal ganglion cells, amacrine and bipolar was not affected by Siah1 overexpression, and there was no significant reduction of proliferating cells in the Siah1 overexpressing retina. We did however detect increased cell death, confirmed by an increase in the number of TUNEL+ cells in the ONL, which was proteasome-dependent, as MG132 treatment rescued the cell death phenotype. Lastly, reduction in rods and cones resulting from increased Siah1 expression was rescued by injection of *cdhr1* mRNA, and to an even greater extent by injection of a Siah1-insensitive *cdhr1a* variant mRNA. Taken together, our work provides the first evidence that Cdhr1a plays a role during early photoreceptor development and that Cdhr1a is regulated by Siah1 via the UPS. This work provides new avenues for investigation into the roles of CDHR1, and now also Siah1, in the predisposition and pathogenesis of inherited cone-rod dystrophy.

## Introduction

According to the World Health Organization (WHO), in 2015 more than 253 million people worldwide were visually impaired, of which 36 million were blind. This number is predicted to increase to 703 million visually impaired people by 2050^1^. Retinal congenital disease is a major contributor to blindness disorders, affecting 4.5 million people worldwide. Congenital retinal blindness is known to be associated with mutations in over 280 different genes^2–4^. While these mutations encompass various portions of the eye, aberrant development and improper maintenance of the retina are the major causes of inherited loss of blindness.

The retina, an extension of the central nervous system, is responsible for not only detecting incoming light, via photoreceptor cells, but also ultimately conveying that signal through the optic nerve and to the brain to be interpreted as vision^5^. Retinal structure and development are fairly well conserved across vertebrates from human to mouse and zebrafish^6^. There are 7 cell types within the retina, which populate 3 nuclear layers. Differentiation of the neural retina generally begins with the innermost neurons of the retina: ganglion cells within the ganglion cell layer (GCL) closely followed by or in parallel with amacrine, bipolar, and horizontal cells of the inner nuclear layer (INL). Müller glia are the last cells of the INL to differentiate^7^. The final retinal cells to differentiate are photoreceptor cells, rods and cones, which are responsible for distinguishing light and dark as well as color, respectively^8^. Photoreceptors populate the outer nuclear layer (ONL) and play an imperative role in detecting light using the outermost portion of the cell, the outer segment^9^. Outer segments are comprised of hundreds of stacked disks which contain the molecular machinery to detect and process light signals via phototransduction^10^. Phototransduction is a highly metabolically demanding process which produces toxic photo-oxidative compounds, requiring outer segments to shed after a period of time and be replenished to maintain proper cell length^11^. Although photoreceptors are imperative for visual system operation, the function of many genes hypothesized to play a role in their development and maintenance have yet to be tested *in vivo*. Mutations in these various genes can lead to the development of a wide spectrum of sight-threatening diseases, such as the commonly known cone-rod dystrophies.

Cone-rod dystrophies are a group of inherited retinal diseases that first affect cone photoreceptors, then rod photoreceptors, or in some cases they are affected simultaneously^12^. Generally, the photoreceptors begin to degenerate, causing progressive loss in visual acuity, color and central vision, and light sensitivity^13^. In order to develop therapeutics for cone-rod dystrophies, understanding the currently unknown mechanism as to how of each of the over 30 genes^14^ implicated in its onset and progression is imperative. To do this, most studies have aimed to elucidate the role of these genes in retinal development and maintenance in vertebrate models such as mouse and zebrafish^15–19^. A well-established candidate gene associated with cone-rod dystrophy which has yet to be explored in a developmental context is photoreceptor specific cadherin CDHR1.

Several clinical studies^3,20–30^ have described mutations in CDHR1 associated with inherited cone-rod dystrophy. Conserved among vertebrates, CDHR1 belongs to the cadherin superfamily of calcium-dependent cell adhesion molecules but is exclusively expressed in photoreceptors of zebrafish, chickens, mice and humans^31^. CDHR1 encodes an intracellular domain, a transmembrane domain in addition to six cadherin repeats^27^. Previous studies using tomography, electron microscopy and immunohistochemistry have unequivocally defined CDHR1 localization to the base of outer segment of photoreceptor cells^31–33^. Additionally, to further determine its precise location in the junction in between the inner segment (IS) and the outer segment (OS), Burgoyne and collaborators^32^ used nanogold cryo-EM to illustrate that CDHR1 forms fibers connecting immature discs at the base of the outer segment. This group hypothesized that CDHR1 is necessary to stabilize and control the disc evagination process during photoreceptor cell outer segment assembly and/or maintenance^30^. A CDHR1 knockout mouse partially supports this hypothesis as well as the correlation of CDHR1 loss of function and cone-rod dystrophy. CHDR1 knockout mice were born with shorter and disorganized photoreceptor outer segments, followed by a progressive loss of photoreceptors (50%) in the next 6 months of life^33^.

While previous studies of CDHR1 have confirmed its importance for photoreceptor development and homeostasis^25^ we lack any understanding of its regulation during these critical events. Interestingly, we have recently characterized a ubiquitin-proteasomal system (UPS) pathway involved in retinal morphogenesis^34^. We observed that the E3 ligase enzyme, Siah1, was expressed throughout the retina during early development and specifically targeted a transcriptional regulator, Nlz2, for degradation. This process ensured timely and precise fusion of the optic fissure of the early retina. When searching for other targets of this E3 ligase based on its well established degron-motif (Pro-[ARTE]-X-Val-X-Pro), we identified zebrafish Cdhr1a as a potential hit. This suggested to us that Siah is a candidate for regulating the turnover of this protein and therefore controlling its function during photoreceptor development or outer segment maintenance.

In our present study we aimed to investigate the Siah-mediated post-translational regulation of Cdhr1a during development of the zebrafish retina. Taken together, our data indicate that stability of Cdhr1a is necessary for zebrafish photoreceptor development and survival and it is subject to regulation through the UPS by Siah1. In particular, we observe significantly reduced photoreceptor number upon induced expression of Siah, but no significant effects on any other retinal cell type. We show these effects are UPS dependent and can be rescued with a proteasome inhibitor (MG132), with *cdhr1a* mRNA, as well as a Siah1 insensitive Cdhr1a variant. Our work provides an *in-vivo* example of vertebrate photoreceptor cell development modulated by UPS-mediated regulation of a gene known to be associated with inherited cone-rod dystrophy.

## Results

### *Siah* and *cdhr1a* are co-localized in the outer nuclear layer during retinal development

As mentioned above, numerous studies have demonstrated *cdhr1* expression in retinal photoreceptor cells and potentially implicated in photoreceptor development^30,31,33^. In contrast, *siah* gene expression during zebrafish retinal development had yet to be described. As such, we sought to investigate *siah* expression and cellular localization during the later stages of retinal development when photoreceptors are maturing. We carried out a comprehensive expression analysis of both *siah* homologues, *siah1* and *siah2l* during retinal development in zebrafish. Using two color fluorescence whole-mount in situ hybridization (FWISH) we examined simultaneous expression of *siah1* or *siah2l* and *cdhr1a* in the zebrafish retina at 3, 4 and 5 dpf (Fig 1). Starting at 3 dpf we observed co-expression of *cdhr1a* and both *siah1* and *siah2l* specifically in the outer nuclear layer (Fig 1A, D). *Siah* expression was also seen throughout the INL and GCL. This pattern of expression was observed up to and including 5 dpf (Fig 1C, F). Co-expression of *siah1* and *cdhr1a* in the ONL therefore indicates that *siah1* and *cdhr1a* are both present and potentially active during photoreceptor cell maturation. This further suggests that Siah1 may have a functional role in regulating Cdhr1a protein stability in photoreceptor cells.

**Figure 1:**
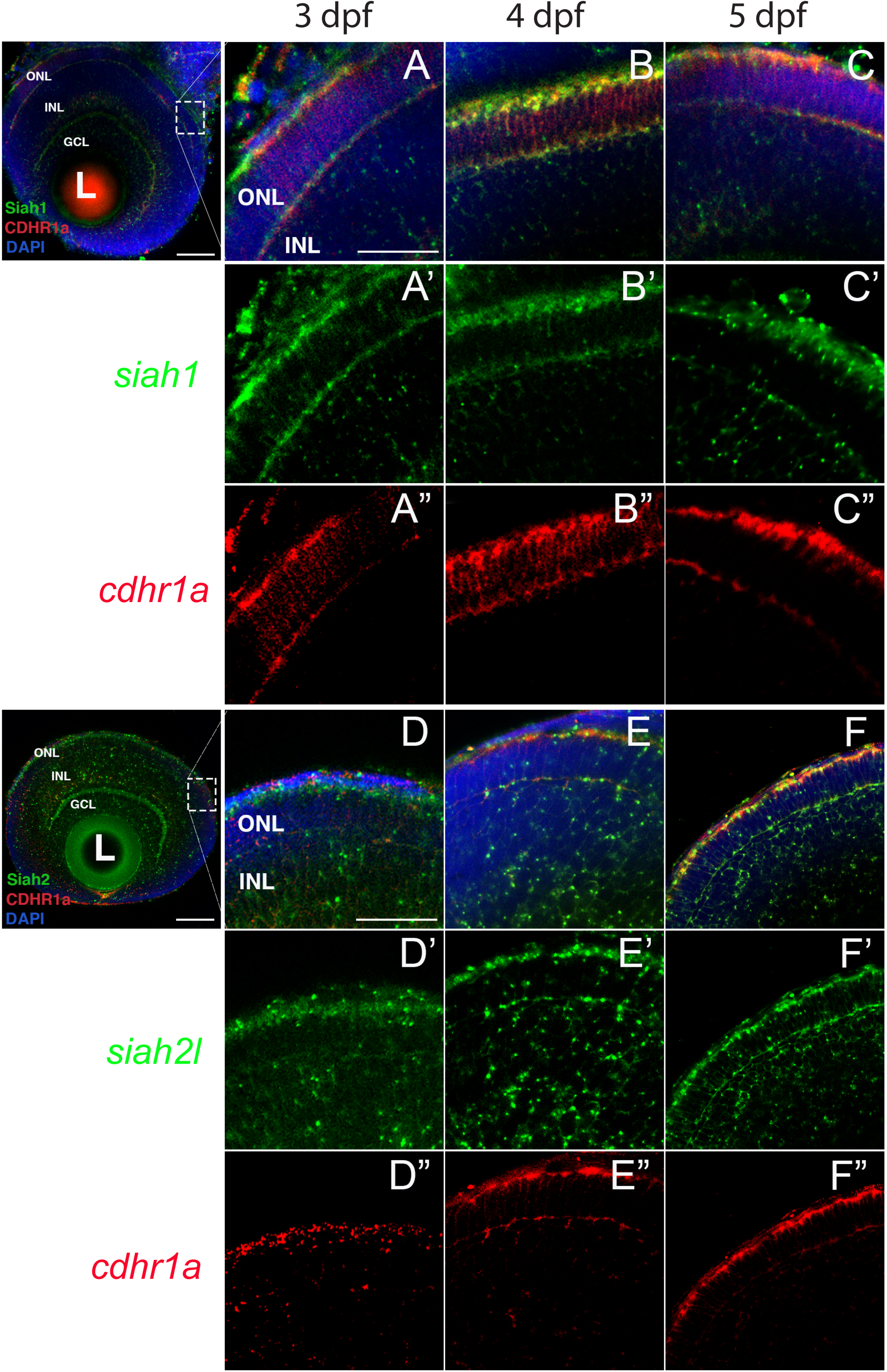
*Siah1* and *cdhr1a* are co-expressed in the outer nuclear layer of the retina. Retinal cryosections from two color fluorescent whole-mount *in situ* hybridization of *siah1* (green) and *cdhr1a* (red) in zebrafish embryos from 3-5 dpf. Magnified images correspond to region of the dashed outline. *Siah1* expression was noted in the outer nuclear and inner nuclear layer from 3-5 dpf (**A’-C’**). *Siah2l* expression was also detected in the outer and inner nuclear layer from 3-5 dpf (**D’-F’**). *cdhr1a* expression was restricted to and throughout only the outer nuclear layer (**A”-C”**). *Siah1* as well as *siah2l* was observed to co-express with *cdhr1a* specifically in the outermost part of the outer nuclear layer (**A-F**). DNA was stained with DAPI (blue). L: lens, ONL: outer nuclear layer, INL: Inner nuclear layer, GCL: ganglion cell layer. Large scale bar = 100μm, small scale bar = 25μm.

### Siah1 localizes to the base of the outer segments in rods and cones

In order to validate our FWISH results, we next examined Siah and Cdhr1a protein localization during photoreceptor maturation. To do so we first obtained zebrafish specific polyclonal antibodies against Siah1 and Cdhr1a. When tested by immunohistochemistry (IHC) in 3-5 dpf retinal sections we observed signal that correlated with our FWISH results. In particular, we observed specific localization of Cdhr1a signal in the outer nuclear layer where the rods and cones reside, while Siah1 signal was detected throughout the retina, including strong signal in the outer nuclear layer (ONL) (Fig 2D-F). To confirm that Siah1 is localized in photoreceptor cells, and to determine to which subcellular region, we performed IHC on retinal sections from transgenic embryos expressing rod and cone reporter constructs, Tg[*XOPS*:GFP] and Tg[*TαC*:eGFP] respectively^35,36^. Our IHC results indicate that Siah1 protein localized to the synaptic terminals of rod (*XOPS*:GFP) and cone (*TαC*:eGFP) photoreceptors, as well as in the connecting cilium from 3 to 5 dpf (Fig 2D’-F’, J’-K’). We observed a similar pattern of localization for Cdhr1a, in particular at the connecting cilium of rods (*XOPS*:GFP) and cones (*TαC*:eGFP) (Fig 2A’-C’, G’-H’). Cdhr1a localization to the primary cilium began at 3 dpf, with low signal, and increased progressively until 5 dpf. In addition to the ONL, Siah1 protein staining was also observed in the inner nuclear layer (INL) and ganglion cell layer (GCL) during the period analyzed. This again corelated with our FWISH data. Collectively, our analysis of mRNA and protein localization for Siah and Cdhr1a indicated that both proteins are expressed in photoreceptor cells and may functionally interact.

**Figure 2:**
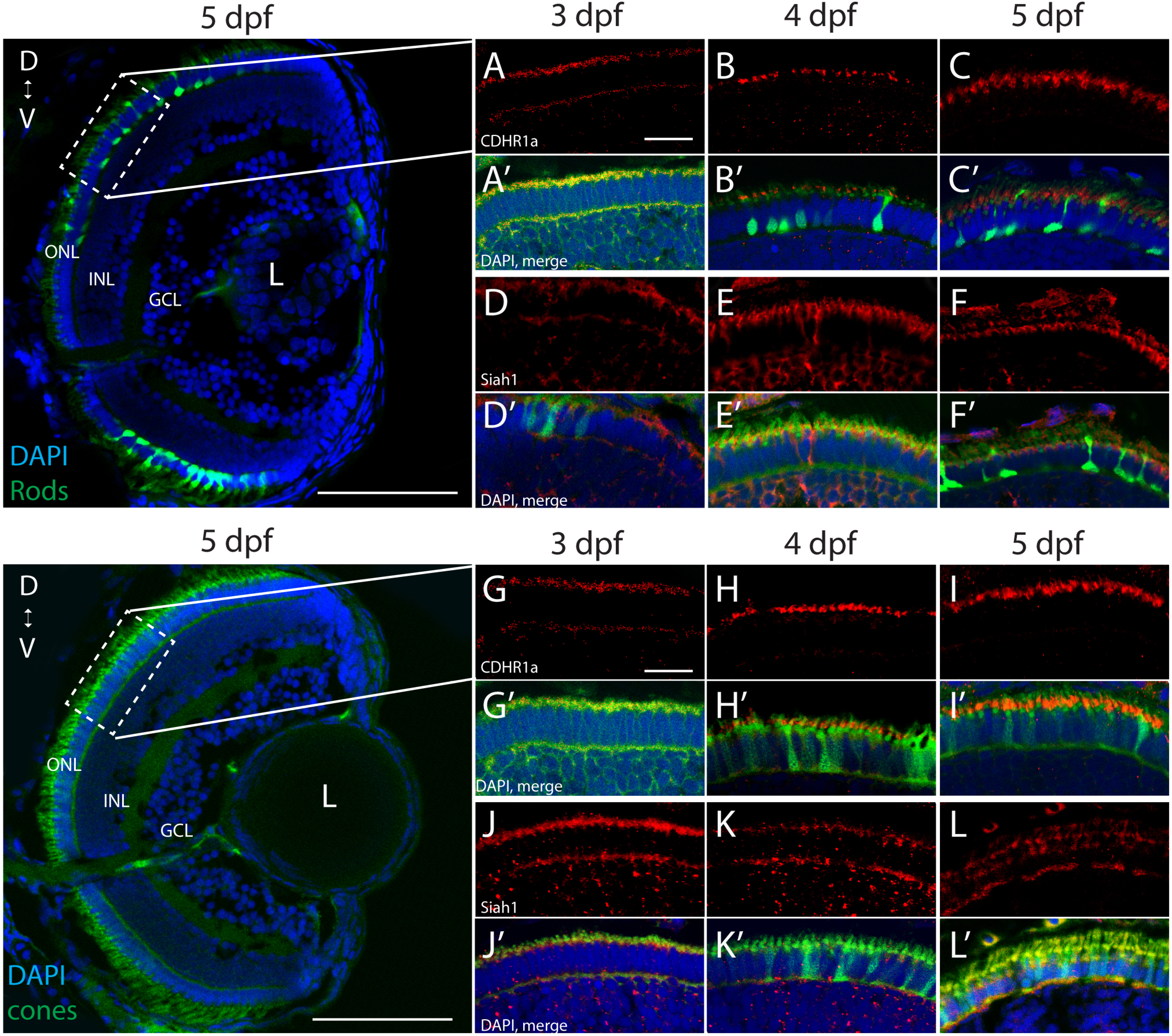
Siah1 and Cdhr1a localize to the photoreceptor primary cilium. Siah1 and Cdhr1a protein localization was determined using IHC in 3-5dpf old Tg[*XOPS*:GFP] or Tg[*TαC*:GFP] retinal cryosections. Cdhr1a signal (red) was detected in the ONL (**A-C, G-I**) and within rod photoreceptors (green) marked by *XOPS*:GFP expression between 3-5dpf (**A’-C’**). Siah1 signal was also detected in the ONL (**D-E, J-L**) and within rod photoreceptors (**D’-E’**). Cdhr1a signal (red) was detected within cone photoreceptors (green) marked by *TαC*:GFP expression between 3-5dpf (**J’-L’**). Siah1 signal was also detected within cone photoreceptors (**D’-E’**). Both Cdhr1a and Siah1 localization within photoreceptors was strongest at the junction of the inner and outer segments. DNA was stained with DAPI (blue). L: lens, ONL: outer nuclear layer, INL: Inner nuclear layer, GCL: ganglion cell layer. D: Dorsal and V: Ventral. Large scale bar = 100μm, small scale bar = 10μm.

### Siah1 targets Cdhr1a for proteasomal mediated degradation

Siah E3 ubiquitin ligase interaction with target proteins is a well-characterized process and it involves interaction through an evolutionarily conserved amino acid motif termed a degron. Zebrafish Cdhr1a protein encodes a Siah degron starting in the 857^th^ amino acid. In vertebrates, CDHR1 is highly conserved, with similarity ranging around 60% when comparing human to zebrafish Cdhr1a (Fig 3A). This includes the degron motif, suggesting that this conserved feature plays an important role in the regulation of Cdhr1a through the UPS. To examine weather Siah regulates Cdhr1a protein degradation, we transiently co-transfected HEK 293T cells with Cdhr1a-FLAG and GFP (control) or Cdhr1a-FLAG and Siah1-myc. Protein levels were subsequently determined by Western blot. As shown in Figure 3B, lysates from cells co-transfected with Cdhr1a-FLAG and Siah1-myc, completely lacked Cdhr1a-FLAG signal while the control co-transfection resulted in the presence of a strong Cdhr1a-FLAG band (Fig 3B), indicating that Cdhr1a is targeted for degradation in the presence of Siah1. To demonstrate a direct effect of Siah1 E3 ligase activity we also co-transfected Cdhr1a-FLAG with an inactivated Siah1 construct (SiahΔRING-myc) which is missing the RING domain and therefore cannot perform the E3-mediated ubiquitin transfer onto its targets. Cell lysate from Cdhr1a-FLAG and Siah1ΔRING-myc co-transfection also contained a strong Cdhr1a-FLAG band. Furthermore, inhibition of the proteasome using MG132 treatment resulted in the retention Cdhr1a-FLAG signal compared to no treatment (Fig 3B). These results demonstrated that Siah1 is directly responsible for the loss of Cdhr1a-FLAG due to proteasomal degradation. To determine whether Cdhr1a targeting by Siah1 requires the degron motif, we constructed a Cdhr1a variant in which the VmP motif of the degron sequence was altered to LmA (cdhr1a^LmA^). The mutations had no effect on the expression of Chdr1a-FLAG and as shown in Figure 3C, Cdhr1a^LmA^-FLAG was completely insensitive to the effects of Siah1-myc. Finally, using co-immunoprecipitation (co-IP) we showed that Siah1-myc, or Siah1ΔR-myc, can be pulled down by Cdhr1a-FLAG (Fig 3D). To ensure the specificity of our co-IP, cells were transfected with Siah1ΔRING-myc alone, showing no pull-down with FLAG antibodies after the co-IP (Fig. 3D). Taken together, these results strongly suggest that Siah1 directly targets Cdhr1a for proteasomal mediated degradation through the degron motif found in Cdhr1a. In light of our findings, we next sought to determine whether Siah-mediated regulation of Cdhr1a protein stability plays a role in zebrafish photoreceptor development.

**Figure 3:**
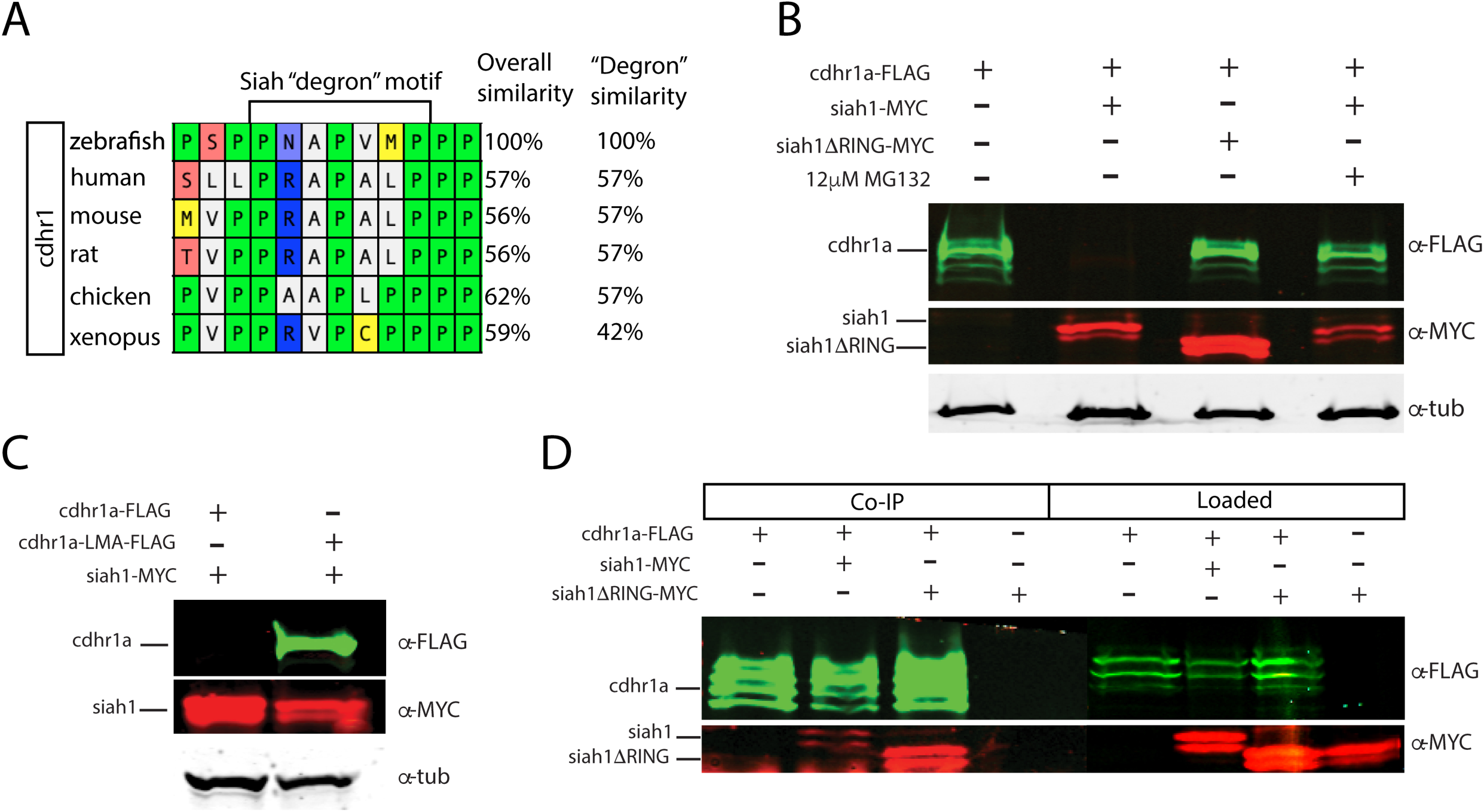
Siah1 targets Cdhr1a for proteasomal degradation. Alignment of CDHR1 degron motif sequence from different vertebrates: Xenopus, chicken, mouse, rat, human and zebrafish outlining overall protein sequence as well as motif conservation (**A**). Western blot analysis of cdhr1a protein stability in response to Siah activity. cdhr1a-FLAG signal is significantly decreased by co-transfection of siah1-myc, but not siah1ΔR-myc or upon MG132 treatment. Alpha/betta tubulin was used as a loading control (**B**). Co-immunoprecipitation of cdhr1a-FLAG co-transfected with siah1-myc or siah1ΔR-myc probed for FLAG (green), MYC (red). Cdhr1a-FLAG is able to pull down both siah1 and siah1ΔRING. (**C**). Western blot analysis of Siah1 targeting specificity. cdhr1a-FLAG signal is significantly decreased by co-transfection of siah1-myc. Signal of cdhr1a^LMA^-FLAG, a cdhr1a variant encoding a non-recognized degron motif, does not decrease upon co-transfection of siah1-myc. Alpha/betta tubulin was used as a loading control (**D**).

### Misregulation of Siah1 activity leads to reduced numbers of photoreceptors

Based on our characterization of *siah1* and *cdhr1a* expression and co-localization in photoreceptor cells, specifically the connecting cilium, and our confirmation that Siah1 targets Cdhr1a for degradation in vitro, we next wanted to determine whether this interaction plays a functional role during photoreceptor cell development. In order to overexpress Siah1 during retinal development, we generated two zebrafish transgenic lines in which Siah1 or the inactive Siah1ΔRING were placed under the control of the heat shock inducible *hsp70* promoter (Fig. 4A). The corresponding lines, Tg[*hsp70*:siah1] and Tg[*hsp70*:siah1ΔRING], were bred to homozygosity. We designed an experimental heat shock approach that would induce Siah1 expression during the developmental window of photoreceptor genesis, between 48 and 72 hpf. To ensure continuous activity of the transgene, we performed the initial head shock at 48 hpf, followed by a repeat heat shock at 60 hpf, and finally fixation at 72 hpf, by which time photoreceptor differentiation is largely completed (Fig. 4B). The efficiency and specificity of the heat shock (HS) system were assessed by whole-mount *in situ* hybridization (WISH) (Fig. 4C). In the absence of elevated temperature examination of *siah1* expression did not suggest any leaky expression from the *hsp70* promoter. Upon heat shock *siah1* and *siah1ΔRING* expression were significantly and ubiquitously elevated (Fig 4C). Tg[*hsp70*:siah1] and Tg[*hsp70*:siah1ΔRING] were next crossed onto the Tg[*XOPS*:GFP] and Tg[*TαC*:eGPP] transgenic lines to assess rod and cone photoreceptor development, respectively. Double transgenic embryos were subjected to the HS protocol. Collected embryos were either imaged whole using confocal microscopy, or cryosectioned for IHC analysis. Confocal imaging of whole-mount embryos clearly indicated a significant decrease in rod and cone cells in Siah1 overexpressing embryos at 72 hpf relative to controls (Fig S1). Quantification of retinal sections confirmed a decrease in the number of rods (Fig. 5A-C) and cones (Fig. 5E-G) in Siah1 HS embryos compared to wild type and siah1ΔRING HS (Fig. 5). The decrease in mature rod photoreceptors in the Tg[*XOPS*:GFP] line was most evident in the ventral portion of the retina (Fig. 5A-C’), where rod photoreceptors initially differentiate^37^ (Fig 5D). Rods in the wildtype HS embryos displayed an elongated cell shape with partially visible outer segments (Fig 5A’) whereas rods in the Siah1 HS embryos appeared wider and without visible outer segments (Fig 5’C’). Rods in Siah1ΔRING HS embryos retained wildtype numbers and morphology (Fig. 5B’, D). Some rods appeared stunted in shape, but most had an elongated structure with visible outer segments, comparable to wildtype HS. When examining Siah1 HS in the *TαC*:eGFP background we observed a decrease in cones both dorsally and ventrally (Fig 5E-G, H). The most striking decrease was again observed in the ventral portion of the retina (Fig. 5G’). Having observed a negative effect of Siah1 overexpression on photoreceptor development, we next sought to determine the extent of these effects in development of other retinal cell types.

**Figure 4:**
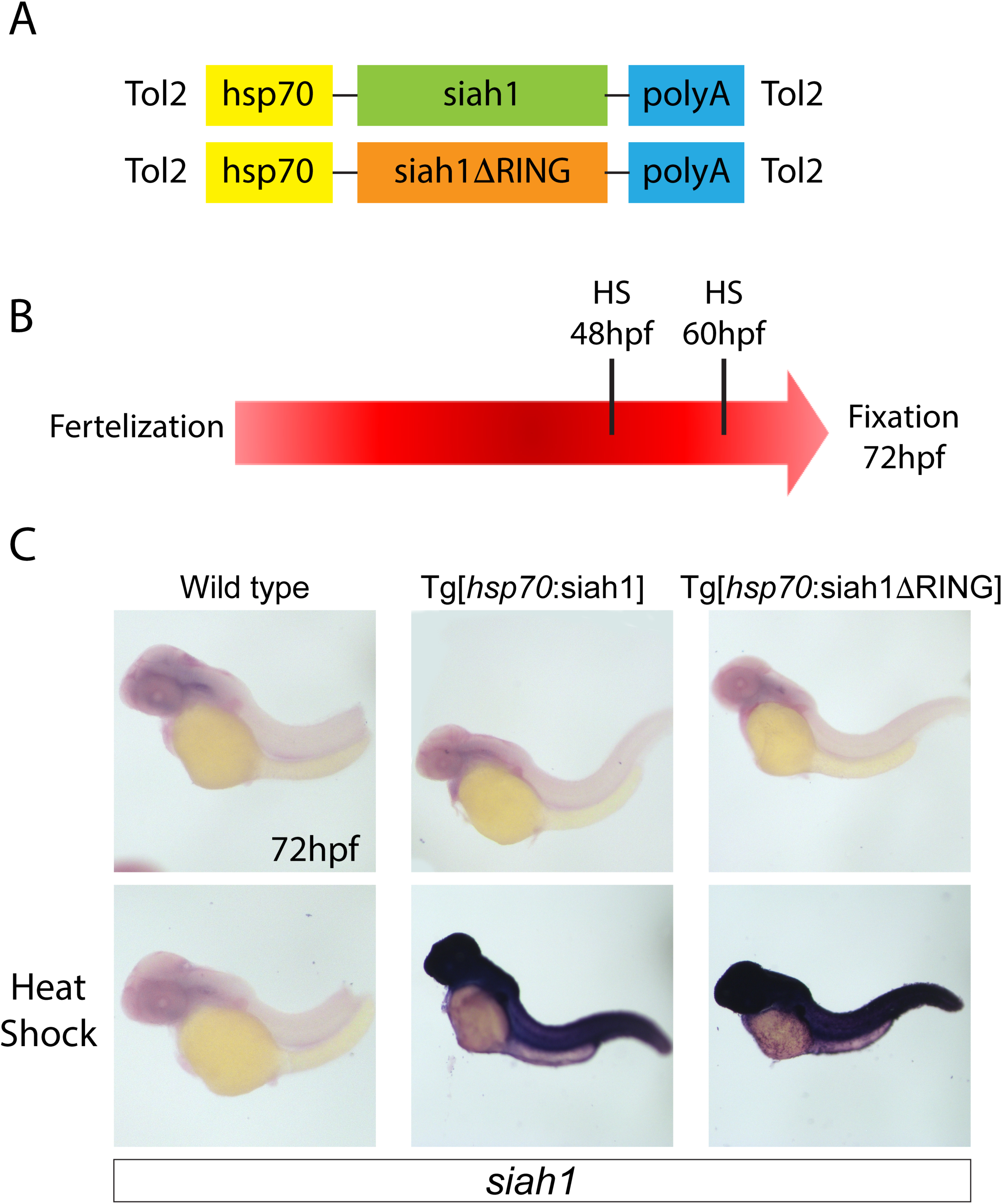
Siah overexpression experimental design. Heat shock line construct schematic (**A**). The experimental design included heat shock for 30 minutes at 48 and subsequently 60 hpf with fixation and analysis at 72 hpf (B). Whole-mount in situ hybridization (WISH) for *siah1* to confirm the effect of heat shock in wild type, Tg[hsp70:Siah1] and Tg[hsp70:Siah1ΔRING] embryos. Heat shock induced a significant increase in *siah1* gene expression in the transgenic lines but not in wild type (**C**).

**Figure 5:**
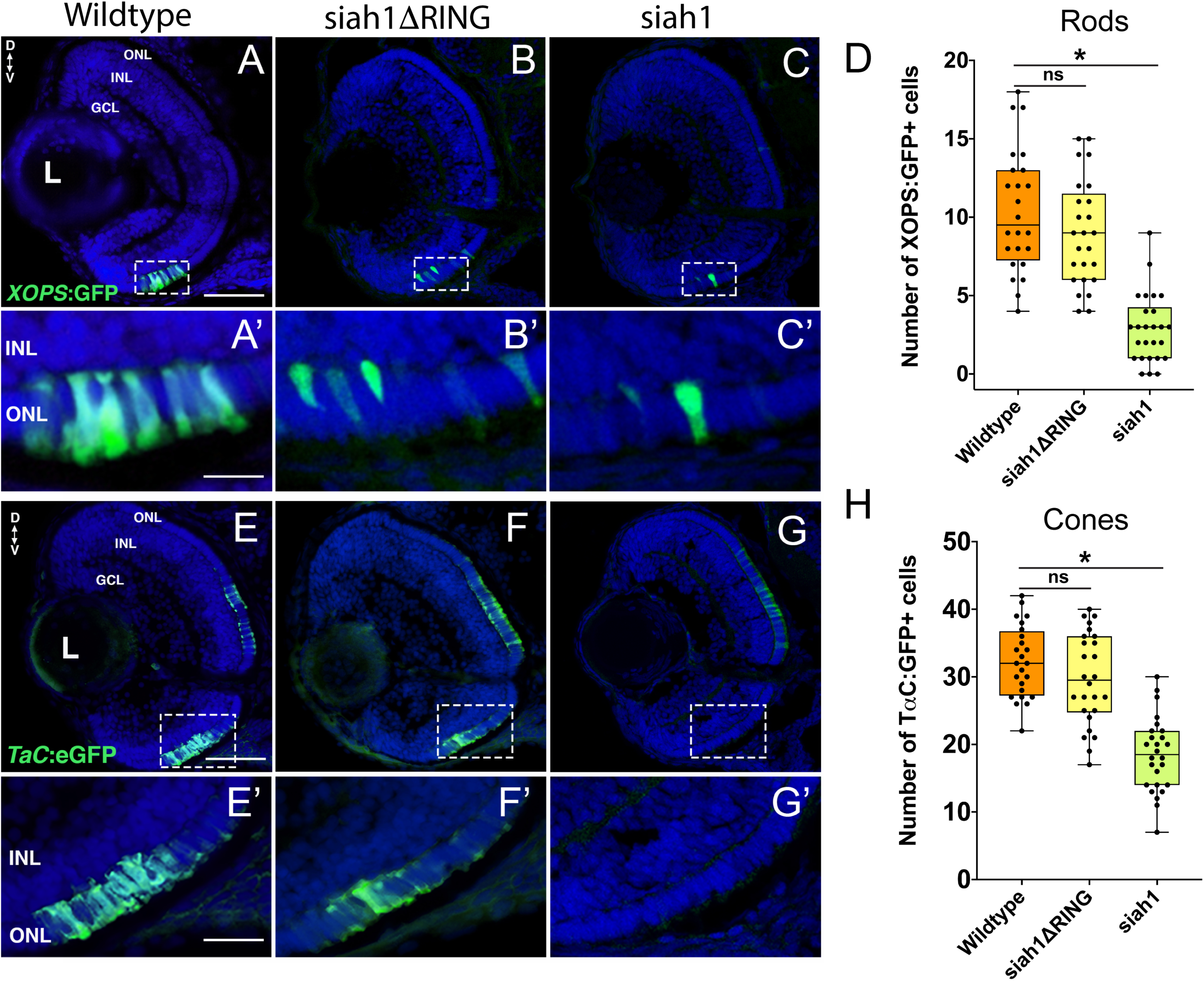
Siah1 overexpression leads to a reduction of rod and cone photoreceptors. Retinal cryosections of Tg[*XOPS*:GFP] (wildtype), Tg[*hsp70*:siah1]/Tg[*XOPS*:GFP] (siah1), and Tg[*hsp70*:siah1ΔRING]/Tg[*XOPS*:GFP], (siah1ΔRING) embryos were analyzed for GFP fluorescence after heat shock (HS) (A-C). The number of rod photoreceptors (green) was significantly decreased in siah1 HS embryos at 72 hpf compared to wildtype and siah1ΔRING (D). Compared to wildtype and siah1ΔRING differentiated rod photoreceptors in Siah1 HS embryos have stunted outer segments (A’-C’). Retinal cryosections of Tg[*TαC*:GFP] (wildtype), Tg[*hsp70*:siah1]/Tg[*TαC*:GFP] (siah1), and Tg[*hsp70*:siah1ΔRING]/Tg[*TαC*:GFP], (siah1ΔRING) embryos were analyzed for GFP fluorescence after heat shock (HS) (E-G). Compared to wildtype and siah1ΔRING, siah1 HS resulted in a significant decrease in the number of cone photoreceptors (green) present the ventral portion of the retina of (E’-G’, H). DNA was stained with DAPI (blue) n=25 embryos, scale bar = 50 μm and 10 μm (A). L: lens, ONL: outer nuclear layer, INL: Inner nuclear layer, GCL: ganglion cell layer, D: Dorsal and V: Ventral.

### Inner retinal neurons are not affected by elevation of Siah1 activity

Since Siah1 can potentially target several proteins and localizes to other regions of the retina (Fig 1), we investigated if overexpression of Siah1 could also impact the development of other cell types in the retina. Ganglion and amacrine cells are among the first retinal neurons to differentiate, a good portion having done so prior to the first HS at 48 hpf^38^. Immunostaining of retinal sections from Siah1 HS embryos with HuC/D indicated that morphology and cell number of ganglion and amacrine cells was unaffected by Siah1 over activation (Fig S2A-D). Similarly, bipolar cells, visualized using PKC*α* immunostaining, were also found to be unaffected by Siah1 overactivation (Fig S2E-H). Horizontal cells, visualized by Prox1 immunostaining, line the outermost part of the INL and have an oblong shape. The Prox1+ horizontal cells within the Siah1 HS embryos had normal morphology (Fig S2E-G) but were decreased in number compared to wildtype and Siah1ΔRING HS (Fig S2H). This phenotype was not as severe as what we observed for rods and cones and was therefore not a focus of our investigation going forward. In summary, from our analysis of ganglion, amacrine, bipolar and horizontal cells, we conclude that the functional consequences of Siah1 overexpression during the 48-72 hpf stage of retinal development is mostly confined to photoreceptors. Our data suggests that high levels of Siah1 activity can specifically alter photoreceptor maturation. As such, we next we began to address the potential mechanisms for how Siah1 activity impacts photoreceptor development.

### Siah1 misexpression does not affect cell proliferation

To determine whether the decrease in rod and cone photoreceptors at 3 dpf in Siah1 HS embryos was due to a delay in differentiation or cell death, we first assessed cell proliferation in the retina. Immunostaining for cells in S phase using PCNA and cells in mitosis with PH3 was conducted to detect potential differences in cell proliferation and cell cycle progression between wildtype, Siah1ΔRING, and Siah1 HS. Our results show no difference in PCNA+ cells in the ciliary marginal zone (CMZ), an area of the retina containing stem and retinal progenitor cells that supports continuous retinal growth^39^, between wildtype, Siah1ΔRING, and Siah1 HS embryos (Fig S3A-C). While photoreceptors do not come from this pool of progenitors, it is an indicator of the rate of cell proliferation in the retina. PCNA was strongly expressed across genotypes and spanned a similar area in the dorsal and ventral portion of the retina. We followed up PCNA analysis with PH3 immunostaining to visualize cells in mitosis, rather than S phase. While there was variation when quantifying PH3+ cells in all genotypes, we found no significant difference in the total number of PH3+ cells when comparing all of our groups (Fig S3G). Interestingly, the distribution of the PH3+ cells did vary across experimental groups (Fig S3H). In the wildtype and Siah1ΔRING HS lines, the majority of PH3+ cells were in the ONL (Fig S3D, E). In contrast, in the Siah1 HS embryos, most PH3+ cells were found in the CMZ and the INL. Taken together, we conclude Siah1 does not regulate cell proliferation or cycle progression, which led us to pursue cell death as a potential mechanism for Siah1 mediated aberrant photoreceptor development.

### Siah1 overexpression results in a proteasome-dependent increase in ONL cell death

We used TUNEL staining to label apoptotic cells in the retina of all genotypes following HS. Additionally, we utilized MG132, which inhibits proteasome activity in order to assay whether the phenotypes observed are dependent on Siah1’s E3 enzymatic activity (control embryos were treated with DMSO). We noted modest cell death, 1-2 cells on average, in the retinas of wildtype and Siah1ΔRING HS embryos (Fig. 6A, C). Apoptotic cells were primarily located in the INL, bordering the CMZ. In contrast, Siah1 HS embryos showed a significant increase in cell death when compared to wildtype and Siah1ΔRING (Fig. 6F). Apoptosis was increased in the GCL, INL, and ONL of the retina (Fig. 6D). To determine whether increased apoptosis was dependent on the proteasome we also treated heat shocked embryos with MG132. Previous work in our lab has shown that 12.5μM of MG132 is effective for embryonic inhibition of the proteasome without toxicity^34^. Cell death was significantly reduced in Siah1 HS embryos treated with MG132, bringing down the average number of TUNEL+ cells to one, which was comparable to wildtype (Fig. 6E, F). After MG132 treatment, any remaining apoptotic cells in Siah1 HS embryos were primarily located in the INL (Fig. 6E). Based on these results, we suspect that reduction of cone and rod photoreceptors upon Siah1 overexpression results from increased cell death of rod and cone progenitors or immature photoreceptors. This effect appears to be dependent on the E3 ligase activity of Siah1 as proteasome inhibition was able to rescue the phenotype. Based on these findings, we next investigated whether Siah1targeting of Cdhr1a contributes to the photoreceptor phenotype of Siah1 overexpressing retinas.

**Figure 6:**
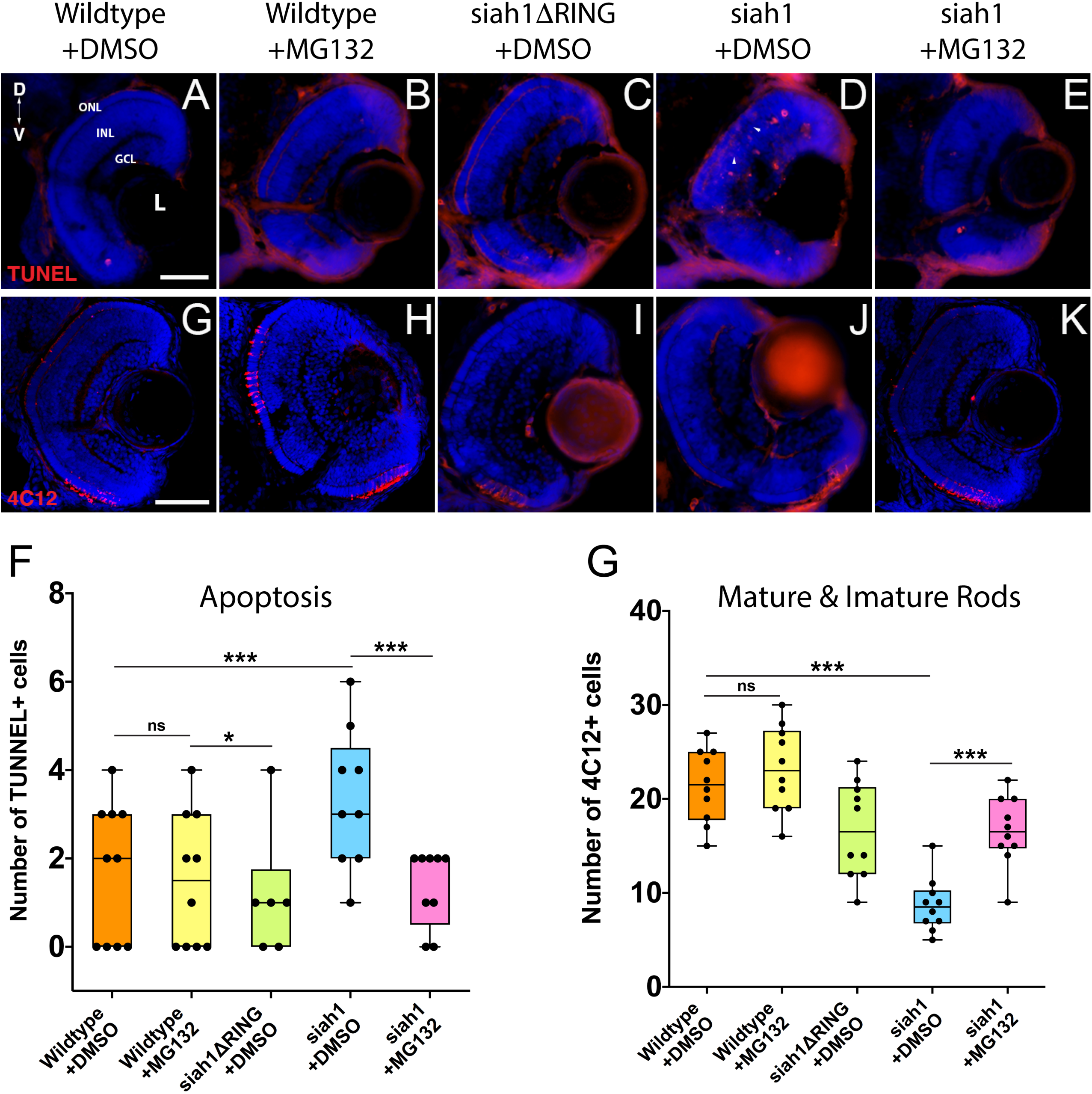
Proteasome inhibition can rescue the decrease in rod photoreceptors and increased apoptosis resulting from Siah1 overactivation. Retinal cryosections from wildtype, Tg[*hsp70*:siah1] (siah1), and Tg[*hsp70*:siah1ΔRING] (siah1ΔRING) embryos heat shocked (HS) and treated with DMSO or MG132, were analyzed using IHC for cell death using TUNEL staining (**A-E**) as well as mature and immature rods using 4C12 antibodies (**G-K**). Number of TUNEL positive cells measured significantly higher in siah1 HS +DMSO embryos compared to all other treatments (**F**). Treatment with MG132 significantly decreased cell death in siah1 HS embryos compared to DMSO to an average comparable to wildtype (**F**). Numbers of mature and immature rod photoreceptors were significantly decreased in Siah1+ DMSO HS embryos treated with DMSO but not with MG132 (**L**). DNA was stained with DAPI (blue). Scale bar = 50μm. L: lens, ONL: outer nuclear layer, INL: Inner nuclear layer, GCL: ganglion cell layer, D: Dorsal and V: Ventral.

### Siah1 targeting of Cdhr1a for proteasomal degradation regulates photoreceptor development and survival

Having shown that MG132 could rescued Siah1 induced apoptosis in the retina, we next examined whether inhibition of apoptosis by MG132 would also rescue photoreceptor development. Treating Siah1 HS embryos with MG132 for 24 hours not only decreased cell death throughout the retina, but also rescued the number of mature and immature rod photoreceptors (Fig 6G-K). Quantification of immature and mature rods, visualized by 4C12 immunostaining^40^, indicated a significant increase in the number of rod cells upon MG132 treatment (Fig. 6L). Mature and immature rod photoreceptors were present in the dorsal and ventral portion of the retina in Siah1 HS MG132 treated embryos (Fig. 6K). The average number of rod photoreceptors was slightly lower in the Siah1 HS MG132 treated embryos compared to wildtype but increased by over 50% when compared to Siah1 HS DMSO treated embryos (Fig. 6L). We observed similar results when examining consequences of MG132 treatment in Tg[*XOPS*:GFP]/Tg[hsp70:siah1] HS embryos (Sup Fig 4A-E). Importantly, cone photoreceptor numbers were also increased upon MG132 treatment, as observed in Tg[*TαC*:GFP]/Tg[hsp70:siah1] HS embryos (Fig S4F-J). Quantification of cone photoreceptors in these embryos showed MG132 treatment led to significantly more cells compared to DMSO treatment (Fig S4J). We therefore conclude that excess Siah1 E3 ligase activity likely leads to increased apoptosis in the ONL and may drive the reduction of both rod and cone photoreceptors.

Having characterized a clear connection between Siah1 and Cdhr1a, we next examined whether Siah1 targeting of Cdhr1a was responsible for the observed phenotypes. As outlined previously, CDHR1 is known to have an established role in photoreceptor maintenance, in particular turnover of outer segment disks^32^. What is less clear is whether CDHR1 plays a role during development of photoreceptors. As such, we hypothesized that Siah1 targets Cdhr1a and the reduction in Cdhr1a protein levels leads to apoptosis and subsequent reduction of photoreceptor progenitor cells. To test this hypothesis, we injected single cell stage Tg*[hsp70*:siah1] embryos with wildtype (WT) *cdhr1a* mRNA, performed our HS treatment and analyzed rod and cone photoreceptors at 72hpf. Injection of mRNA had no observable effect on WT or Siah1ΔRING HS embryos, however, in Siah1 HS embryos we observed a significant increase in the number of both rod and cone photoreceptors compared to Siah1 HS alone at 72hpf (Fig 7A,B). Both mature and immature rod cells were increased in number when Siah1 HS embryos were injected with WT *cdhr1a* mRNA (Fig 7D). Similar outcomes were observed when using Tg[*XOPS*:GFP]/Tg[*hsp70*:siah1] embryos (Fig 7E-F, H). Not only were the number of cells increased, but the rod cells appeared elongated, contained outer segments and were evenly spaced. When examining cone cells using the Tg[*TaC*:eGFP]/Tg[*hsp70*:siah1] we also documented that injection of WT *cdhr1a* mRNA rescued the number of cone cells to levels comparable to wildtype or siah1ΔRING HS embryos (Fig 8). Our results show that an excess of Cdhr1a can overcome the Siah1 targeting and therefore protect development of photoreceptor cells.

**Figure 7:**
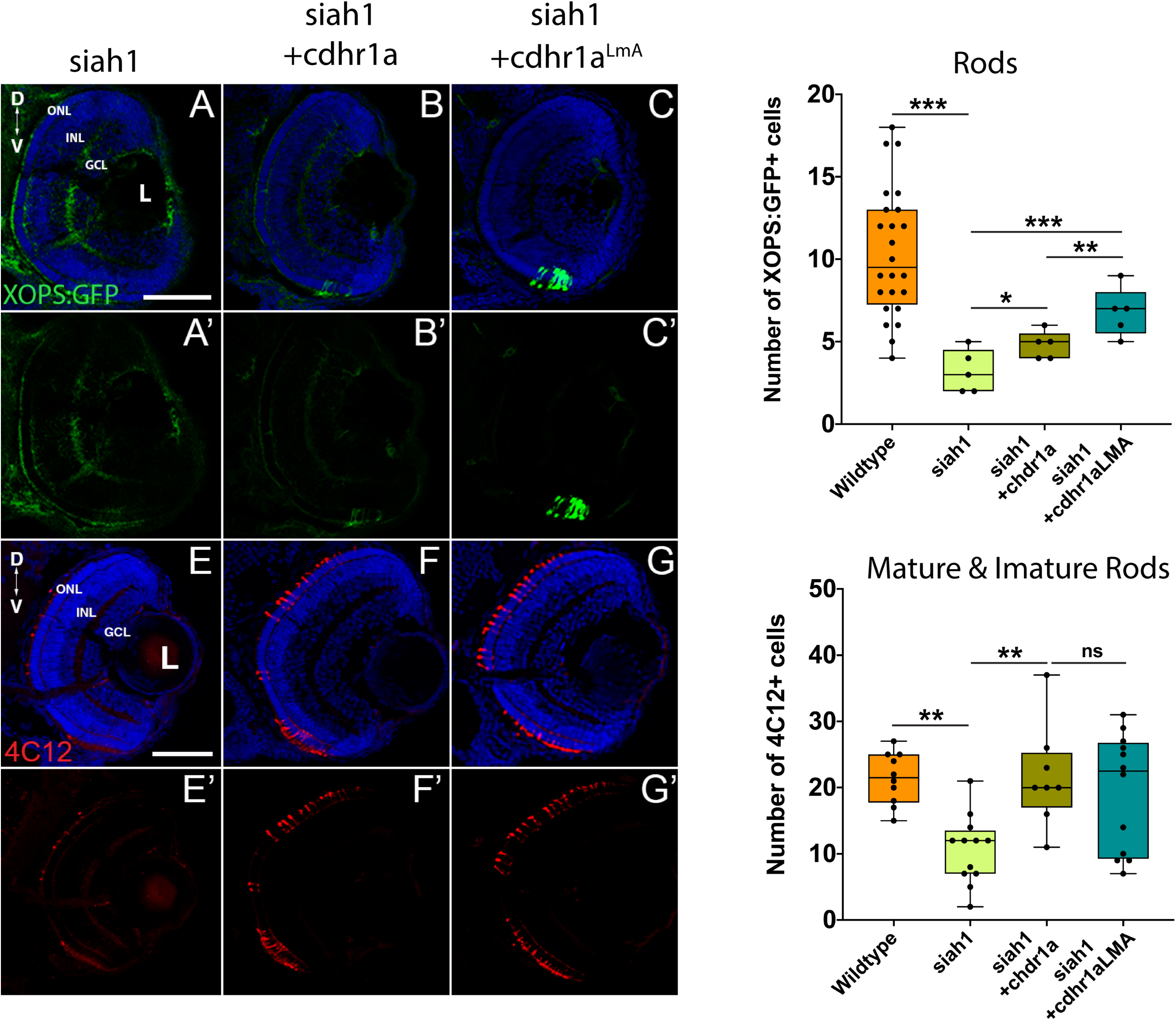
Rod photoreceptor development relies on sufficient levels of Cdhr1a. Retinal cryosections from Tg[*hsp70*:siah1]/Tg[*XOPS*:GFP] (siah1), injected with wildtype *cdhr1a* or *cdhr1a*^*LMA*^ mRNA were heat shocked (HS) and analyzed for immature and mature rod cells using IHC for 4C12 (red) (**A-C’**). Injection of both *cdhr1a* and *cdhr1a*^*LMA*^ mRNA increased the number of immature and mature rod cells compared to siah1 HS alone (**D**). Retinal cryosections from Tg[*hsp70*:siah1]/Tg[*XOPS*:GFP] (siah1), injected with wildtype *cdhr1a* or *cdhr1a*^*LMA*^ mRNA were heat shocked (HS) and analyzed for GFP signal (green) (**E-G’**). Injection of both *cdhr1a* and *cdhr1a*^*LMA*^ mRNA increased the number of GFP+ rod cells compared to siah1 HS alone, with *cdhr1a*^*LMA*^ giving a significantly stronger response (**H**). DNA was stained with DAPI (blue). Scale bar = 50μm. L: lens, ONL: outer nuclear layer, INL: Inner nuclear layer, GCL: ganglion cell layer, D: Dorsal and V: Ventral.

**Figure 8:**
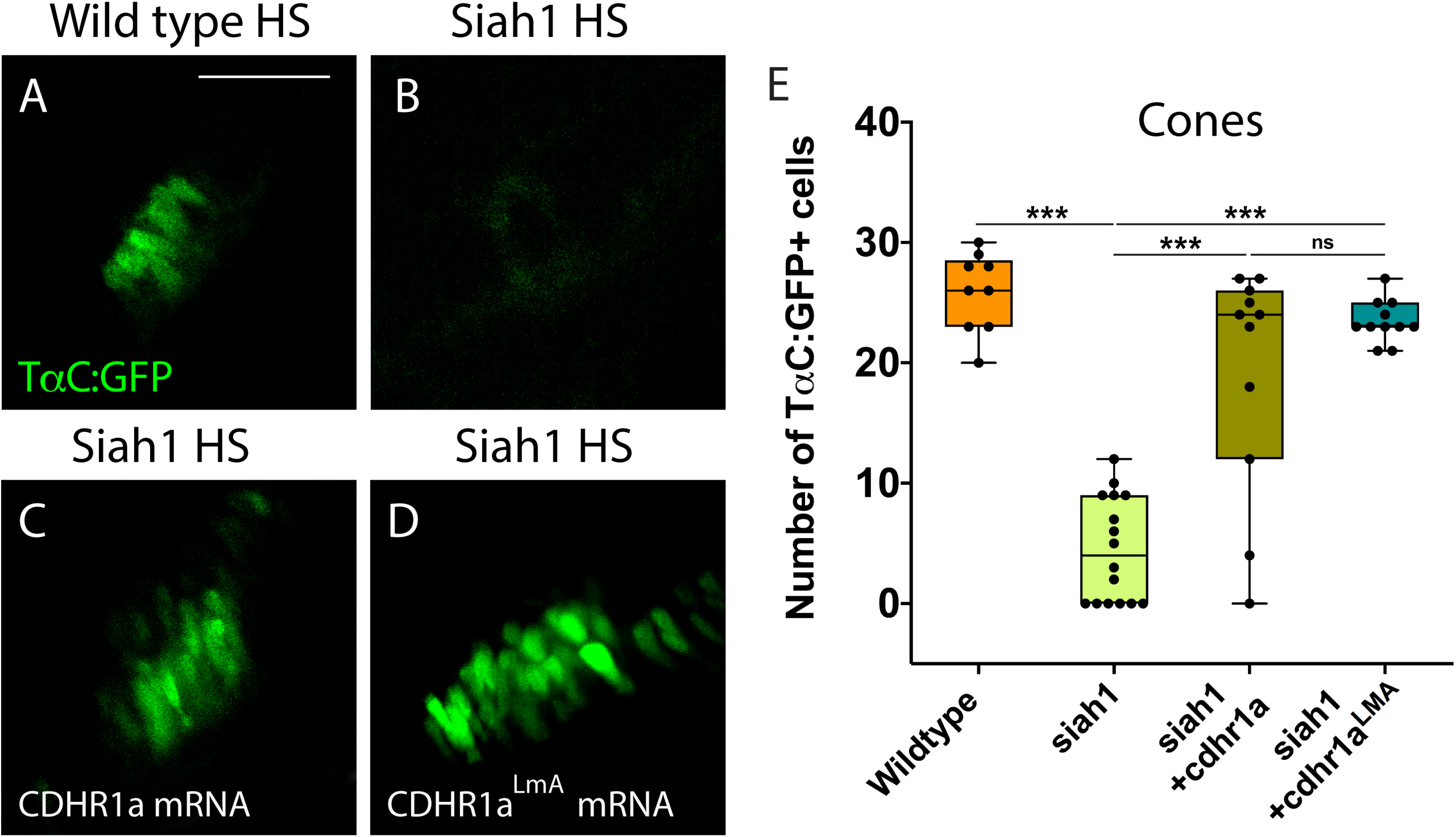
Cone photoreceptor development relies on sufficient levels of Cdhr1a. Confocal stacks of heat shocked (HS) Tg[*TαC*:GFP] (wildtype) and Tg[*hsp70*:siah1]/Tg[*TαC*:GFP] (siah1) embryos or those injected with *cdhr1a* mRNA or *cdhr1a*^*LMA*^ mRNA were analyzed in 3D for GFP fluorescence (**A-D**). Injection of both wildtype and the LMA variant of *cdhr1a* mRNA resulted in numbers of GFP+ cone cells comparable to that of wildtype (**E**). Scale bar = 50 μm.

To further extend our analysis, we also attempted rescue of the Siah1 HS phenotype using the Cdhr1a degron mutant construct, *cdhr1a*^*LmA*^. Based on the insensitivity of Cdhr1a^LmA^ to Siah1 activity we predicted it would have an enhanced rescue effect. Single cell embryos were injected with *cdhr1a*^*LmA*^ mRNA and subjected to our HS protocol. Strikingly, *cdhr1a*^*LmA*^ was more efficient at protecting rod (Fig 7C, G) and cone cells (Fig 8A-D) from the effects of Siah1 overexpression than WT *cdhr1a*. In particular, injection of *cdhr1a*^*LmA*^ resulted in significantly more *XOPS*:GFP+ mature rod photoreceptor cells compared to WT *cdhr1a* (Fig 7D,H). Both WT and LmA were able to rescue the number of *TaC*:eGFP positive cones, but with LmA having a much tighter distribution (Fig 8E). Taken together, we show that deficiencies in rod and cone photoreceptor development upon induction of Siah1 activity correlate with levels of Cdhr1a protein. We therefore conclude that Siah1-mediated regulation of Cdhr1a protein levels is important during photoreceptor development.

## Discussion

Several studies in the past decade have associated mutations in the human CDHR1 gene with cone-rod dystrophies ^3,20–22,24–26,28–30,41,42^. While the mechanism as to how loss of CDHR1 function affects pathogenesis of cone-rod dystrophies is unknown, several studies have reinforced its importance to photoreceptor cell biology by characterizing its protein localization ^31–33^ and necessity for photoreceptor disk renewal ^30^. In particular, it has been shown that CDHR1 links immature disks to the inner segment prior to their incorporation into the outer segment^32^. However, none of the current studies examined in detail whether CDHR1 had any involvement in vertebrate photoreceptor cell development. This is of particular note, especially when considering possible inheritable associations with cone-rod dystrophy predisposition. In this study we describe a post-translational modification mechanism, controlled by the Siah1 E3 ubiquitin ligase, which regulates the stability of Cdhr1a to mediate vertebrate photoreceptor cell maturation, and survival. Our work outlines the first observation of a functional role for Siah1 and Cdhr1a during photoreceptor development, which may in future studies be utilized to examine the mechanism of cone-rod dystrophy pathogenesis.

*Cdhr1* encodes a photoreceptor cell specific cadherin; a single-pass transmembrane glycoprotein with calcium-dependent adhesive abilities as well as signaling functions. Cadherin extracellular domains contain several tandem repeats of negatively charged amino acids which are responsible for interaction with extracellular molecules including other cadherins ^43–45^. During eye development, cadherins have been implicated in the separation of the invaginated lens vesicle from the surface ectoderm ^44^, initiation and elongation of the RGC axons and dendrites ^45–48^, differentiation of RGC and amacrine cells ^43,45^, activation of proliferation in the eye primordium ^45,47,49^ and retinotectal axon projection ^43^. In line with previous studies ^31–33^, we confirmed *cdhr1a* gene expression and protein localization to be specifically in the base of the outer segment of zebrafish rod and cone photoreceptor cells. Zebrafish Cdhr1a protein localized to a narrow stalk region, known as the connecting cilium of the photoreceptor cell. The connecting cilium bridges the outer segment with the cell body and is critical for proper trafficking of proteins, like rhodopsin, from the cell body to the outer segment^50^. This region is also the site of new disk assembly and release during maintenance of rod outer segments. Mutations in structural proteins of this region are known to associate with juvenile Retinitis Pigmentosa^50^, highlighting its relevance in photoreceptor cell development. Furthermore, cadherins, through their cytoplasmic domains are able to link with the cytoskeleton by interacting with catenins^44^. These interactions are responsible for maintaining polarization of the highly stratified epithelial tissues, such as the retina ^43^. When cells of epithelial tissues have blocked their ability to maintain cellular adhesion with the surrounding cells or the extracellular matrix by the loss of cadherin function, for example, they undergo a process of apoptosis called anoikis ^44^.

Our observation that zebrafish Cdhr1a protein localizes to the connecting cilium as soon as photoreceptor cells are formed reinforces the notion of CDHR1’s importance to photoreceptor biology. However, we lacked an understanding of its regulation. Our FWISH results indicated that expression of *siah1* and *siah2l* co-localizes with *cdhr1a* in the ONL from 3 and up to 5 dpf of the zebrafish retina. Furthermore, using a cell culture model, we were able to demonstrate direct interaction, via co-IP experiments, and showed that Siah1 targets Cdhr1a for proteasomal degradation. In addition, we show that inactivation of the Siah1 E3 domain (siah1ΔRING), proteasomal inhibition (MG132), or mutation of the Siah1 degron motif (*chdr1a*^LmA^) prevents Cdhr1a degradation. Taken together, our data strongly supports that Cdhr1a is a direct target for Siah1 and that both are expressed in the same place and at the same time.

To examine the consequences of Siah1-mediated regulation of Chdr1a stability during photoreceptor development we used a heat shock mediated overexpression approach. This enabled us to control the timing and extent of Siah1 overactivation. In particular we wanted to avoid interfering with early embryonic development so as to prevent non-specific phenotypes. Siah1 overexpression resulted in a significant decrease in the amount of rod and cone photoreceptor cells. All other retinal cell types were unaffected by Siah1 overexpression. The reduction in rods and cones coincided with a significant increase in TUNEL+ apoptotic cells in the retina, and in particular in the ONL (Fig. 6D). As expected, inhibition of proteasome activity rescued the Siah1 overactivation phenotype. The cell death we noted could result from cellular loss of contact with the extracellular matrix and/or neighboring cells mediated by Chdr1a in photoreceptor precursor cells. Recently, a mouse conditional double knockout for E and N-cadherin had increased number of TUNEL-positive cells in the lens^44^. Cell death was also noted in the retina of cdh11 and cdh4 morphants^43,45^. Interestingly, when we injected WT or the LmA Siah insensitive *cdhr1a* mRNA and overactivated Siah1, rod and cone photoreceptor numbers were rescued. We therefore propose that Cdhr1a stability in the presence of excess Siah1 is critical for photoreceptor cell survival during development. This is supported by the fact that overactivation of Siah1, which will lead to a decrease in Chdr1a protein levels, leads to increased apoptosis and ultimately significant reduction in rods and cones. In addition, preliminary results from Alt-R-CRISPR injections, which have been shown to be highly efficient in generating biallelic indel mutations and therefore enable examination of phenotypes in F0s, indicate that loss of Cdhr1a function also leads to a decrease in rods and cones (data not shown).

When comparing our results to those of previous studies^31–33^, we propose three potential roles for Cdhr1a during photoreceptor development. First, based on its cadherin function, Cdhr1a may be required for the organization of cytoskeletal elements at the base of newly forming outer segments. In its absence, failure of outer segment formation may trigger apoptosis and subsequent reduction in photoreceptor cells. Second, Cdhr1a, via its extracellular domains, could interact with extracellular matrix in the ONL. In this proposed role, Cdhr1a contributes to either photoreceptor precursor migration and targeting or subsequent photoreceptor adhesion required for survival. The absence of Cdhr1a function could therefore either reduce the number of photoreceptor precursors reaching the ONL leading to reduction of mature rods and cones or may affect maturing rod and cone survival due to absent or improper cell-cell adhesion. Both of our hypotheses are supported by results from our Siah overexpression experiments, which lead to a reduction in Cdhr1a function. Furthermore, our model finds clear support from the transcriptomic analysis recently employed by Kaewkhaw and collaborators^51^ showing increased levels of CDHR1 in photoreceptor progenitor cells during differentiation in 3D human retina cultures. Lastly, based on its localization, Cdhr1a is predicted to regulate the release of newly formed outer segment disks to ensure proper function of rods and cones^32^. One could imagine that assembly of the very first disks would also require Cdhr1a function and in its absence this process might fail and lead to apoptosis. Upon decrease of Cdhr1a levels, the connection between the innermost outer segment disks and the inner segment of the photoreceptor could weaken, preventing outer segment disk formation and eventual death of the photoreceptor prior to maturity (Fig. 8). Interestingly, it currently remains unknown as to how Cdhr1a releases the disks. It may be possible that Siah1 targets Cdhr1a for degradation and this regulates the timely release of the disks. Investigating Siah1 function in juvenile and adult photoreceptors will need to be performed to assess these possibilities.

In conclusion, we provide the first direct evidence that Cdhr1a plays a critical role during photoreceptor development, maturation, and survival. Furthermore, we show that Cdhr1a is directly regulated by the UPS via interaction with Siah1. Our findings have new implications for examination of Cdhr1a-associated cone-rod dystrophy as well as the role of UPS during photoreceptor development. Future studies will focus on the exact mechanism of Cdhr1a function in both photoreceptor progenitors as well as immature rods and cones. Furthermore, it will be important to assess the role of Cdrh1a and Siah1 during retinal regeneration and adult photoreceptor outer segment maintenance. Understanding these mechanisms will be imperative to identifying therapeutic strategies for the growing population of individuals suffering from sight-threatening diseases such as cone-rod dystrophy.

---

**Supplemental Figure 1:**
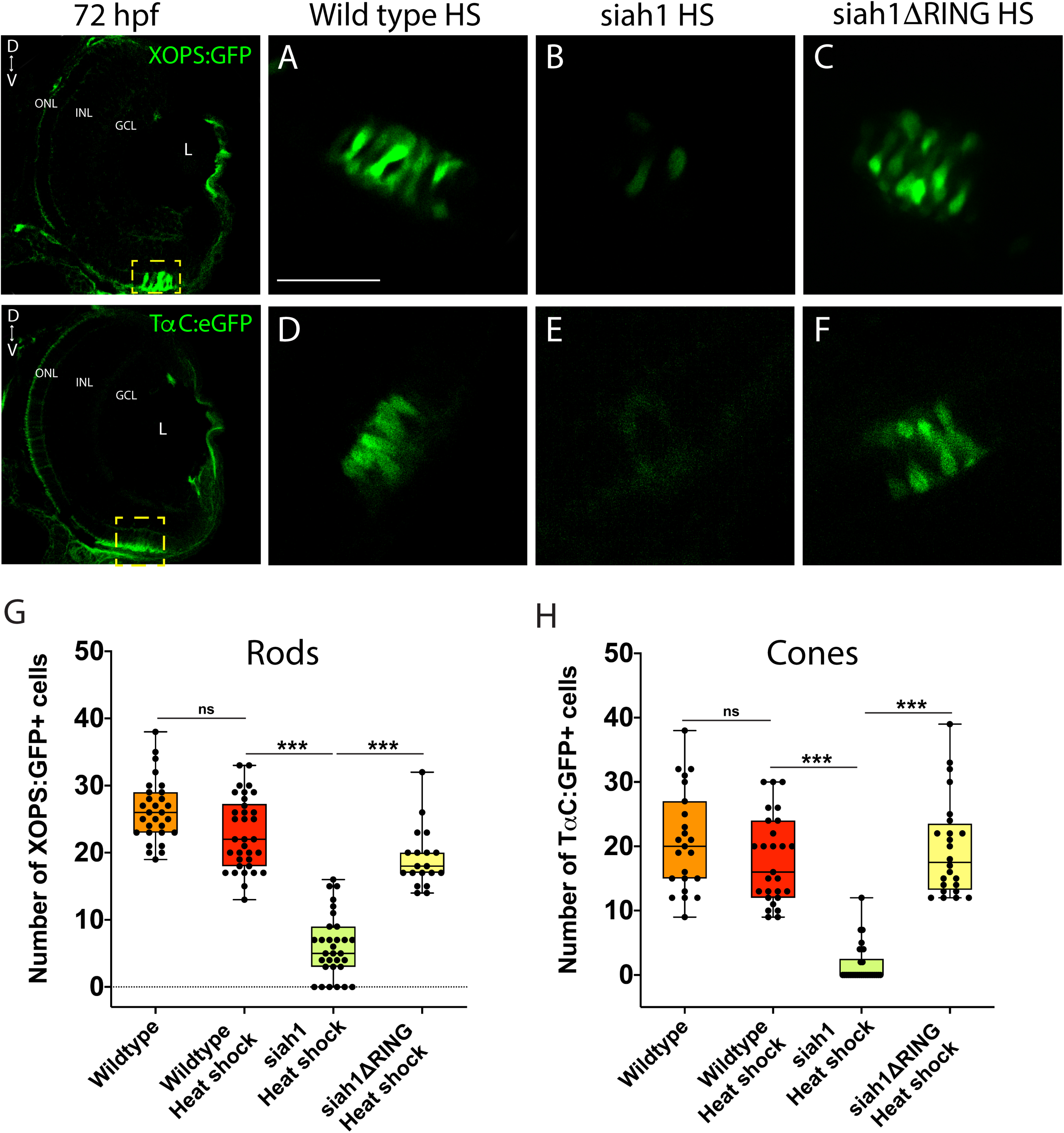
Siah1 overexpression decreases the number of rods and cones. Confocal stacks of heat shocked (HS) Tg[*XOPS*:GFP] (wildtype), Tg[*hsp70*:siah1]/Tg[*XOPS*:GFP] (siah1), and Tg[*hsp70*:siah1ΔRING]/Tg[*XOPS*:GFP], (siah1ΔRING) embryos were collected analyzed in 3D for GFP fluorescence (**A-C**). Region analyzed and presented is outlined in yellow. Overexpression of Siah1 resulted in significantly fewer GFP+ rod cells (**G**). Confocal stacks of heat shocked (HS) Tg[*TαC*:GFP] (wildtype), Tg[*hsp70*:siah1]/Tg[*TαC*:GFP] (siah1), and Tg[*hsp70*:siah1ΔRING]/Tg[*TαC*:GFP], (siah1ΔRING) embryos were analyzed in 3D for GFP fluorescence (**D-F**). Region analyzed and presented is outlined in yellow. Overexpression of Siah1 resulted in significantly fewer GFP+ cone cells (**H**). Scale bar = 50μm.

**Supplemental Figure 2:**
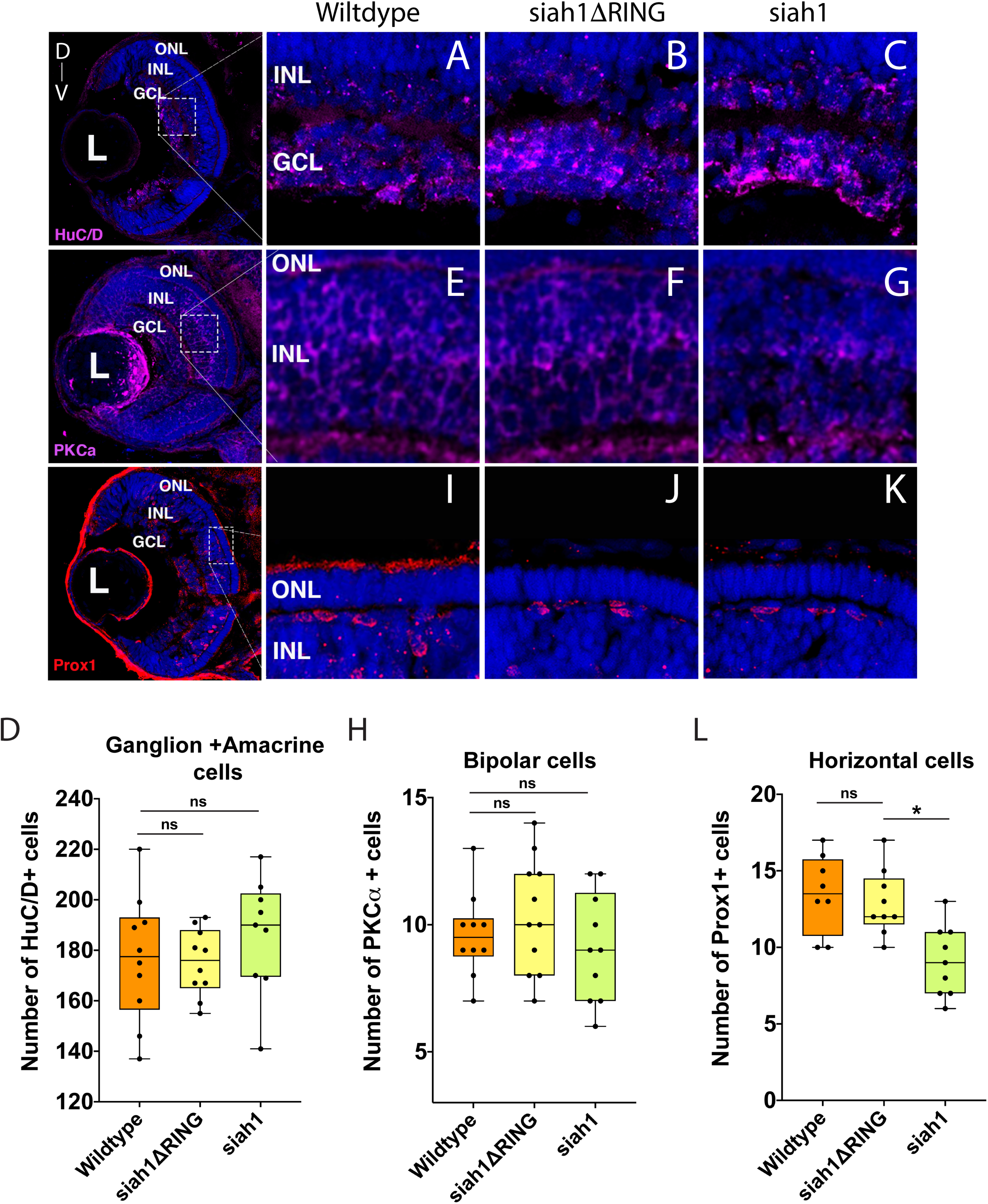
Inner retinal neurons are not affected by Siah1 overexpression. Retinal cryosections from wildtype, Tg[*hsp70*:siah1] (siah1), and Tg[*hsp70*:siah1ΔRING] (siah1ΔRING) embryos heat shocked (HS) and analyzed for effects on retinal inner neurons using IHC. Retinal ganglion and amacrine cells were visualized and quantified using Huc/D staining (**A-D**). Bipolar cells were visualized and quantified using PKC*α* (**E-H**). Horizontal cells were observed and quantified using prox1 staining (**I-L**). DNA was stained with DAPI (blue). Scale bar = 50μm. L: lens, ONL: outer nuclear layer, INL: Inner nuclear layer, GCL: ganglion cell layer, D: Dorsal and V: Ventral.

**Supplemental Figure 3:**
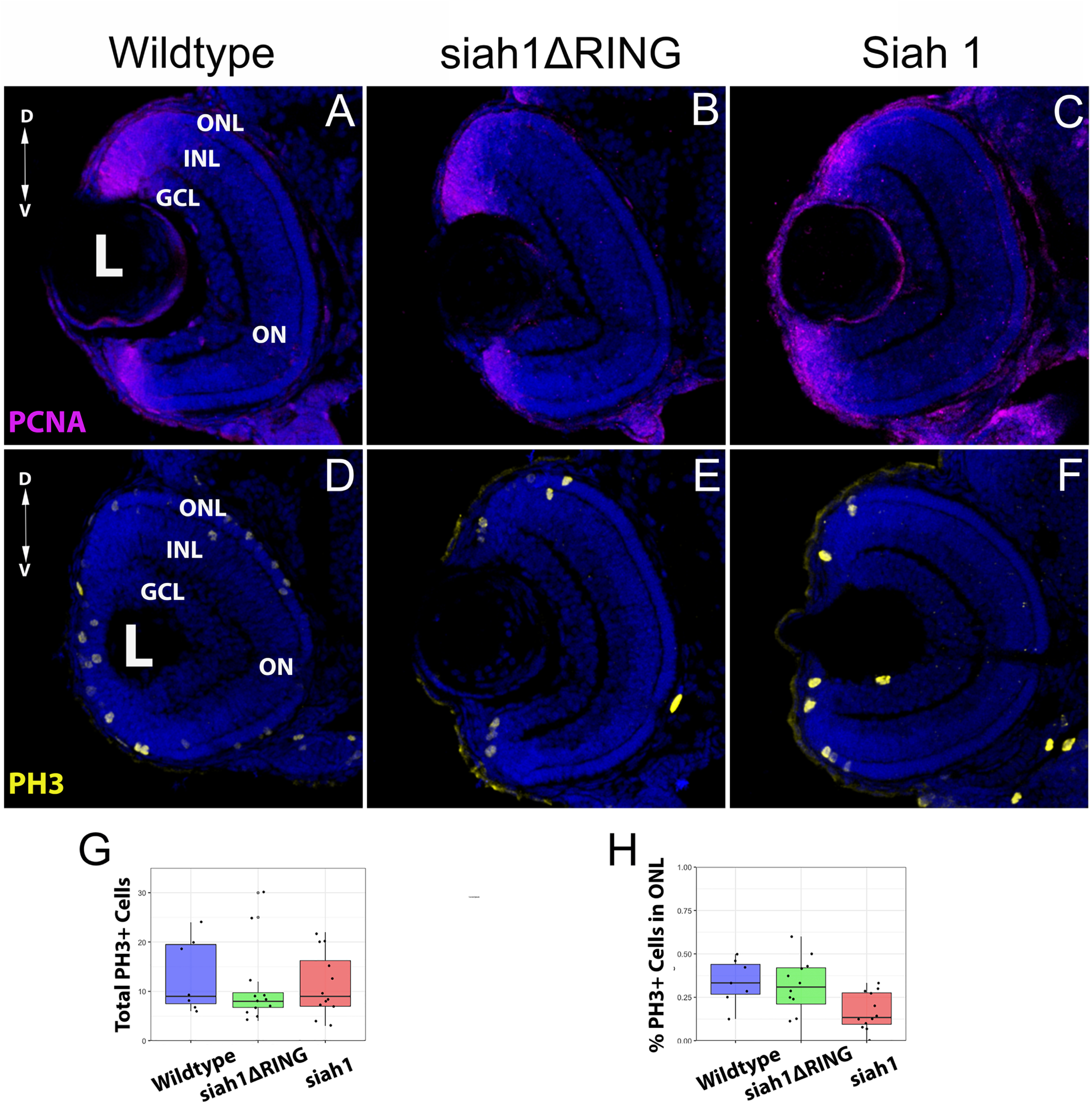
Siah1 does not affect retinal cell proliferation or cell cycle progression. Retinal cryosections from wildtype, Tg[*hsp70*:siah1] (siah1), and Tg[*hsp70*:siah1ΔRING] (siah1ΔRING) embryos heat shocked (HS) and analyzed for cell cycle status using PCNA (**A-C**) and PH3 (**D-F**) IHC staining. Number of PCNA or PH3 positive cells did not significantly change upon Siah1 overexpression (**G-H**). DNA was stained with DAPI (blue). Scale bar = 50 μm. L: lens, ONL: outer nuclear layer, INL: Inner nuclear layer, GCL: ganglion cell layer, D: Dorsal and V: Ventral.

**Supplemental Figure 4:**
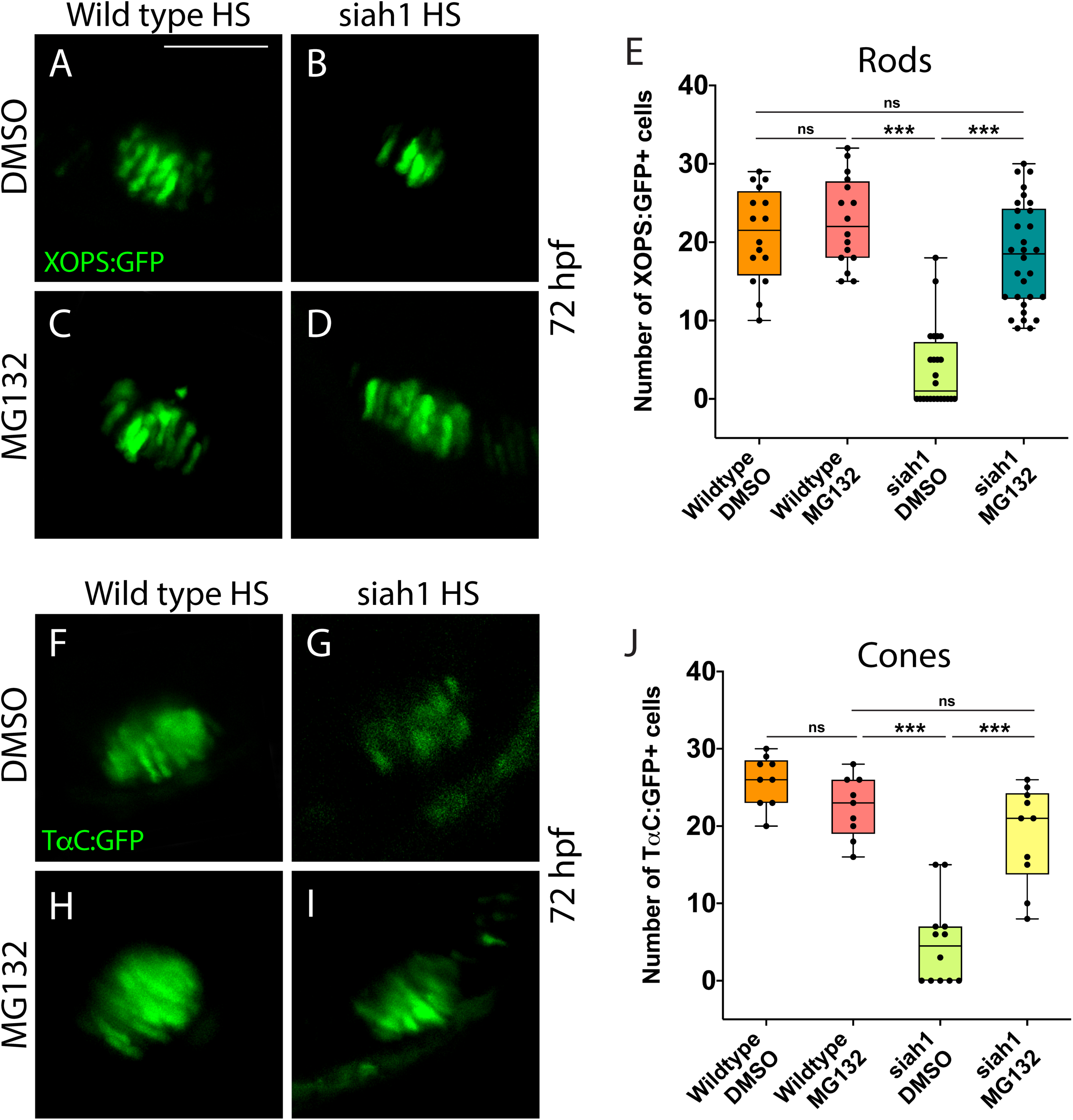
Proteasome inhibition rescues Siah1 overexpression phenotype. Confocal stacks of heat shocked (HS) Tg[*XOPS*:GFP] (wildtype), Tg[*hsp70*:siah1]/Tg[*XOPS*:GFP] (siah1), and Tg[*hsp70*:siah1ΔRING]/Tg[*XOPS*:GFP], (siah1ΔRING) embryos treated with DMSO or MG132 were collected analyzed in 3D for GFP fluorescence (**A-D**). Treatment with MG132 prevented a significant decrease in GFP+ rod cells compared to DMSO in siah1 HS embryos (**E**). Confocal stacks of heat shocked (HS) Tg[*TαC*:GFP] (wildtype), Tg[*hsp70*:siah1]/Tg[*TαC*:GFP] (siah1), and Tg[*hsp70*:siah1ΔRING]/Tg[*TαC*:GFP], (siah1ΔRING) embryos treated with DMSO or MG132 were analyzed in 3D for GFP fluorescence (**F-I**). Treatment with MG132 prevented a significant decrease in GFP+ cone cells compared to DMSO in siah1 HS embryos (**J**). Scale bar = 50 μm.

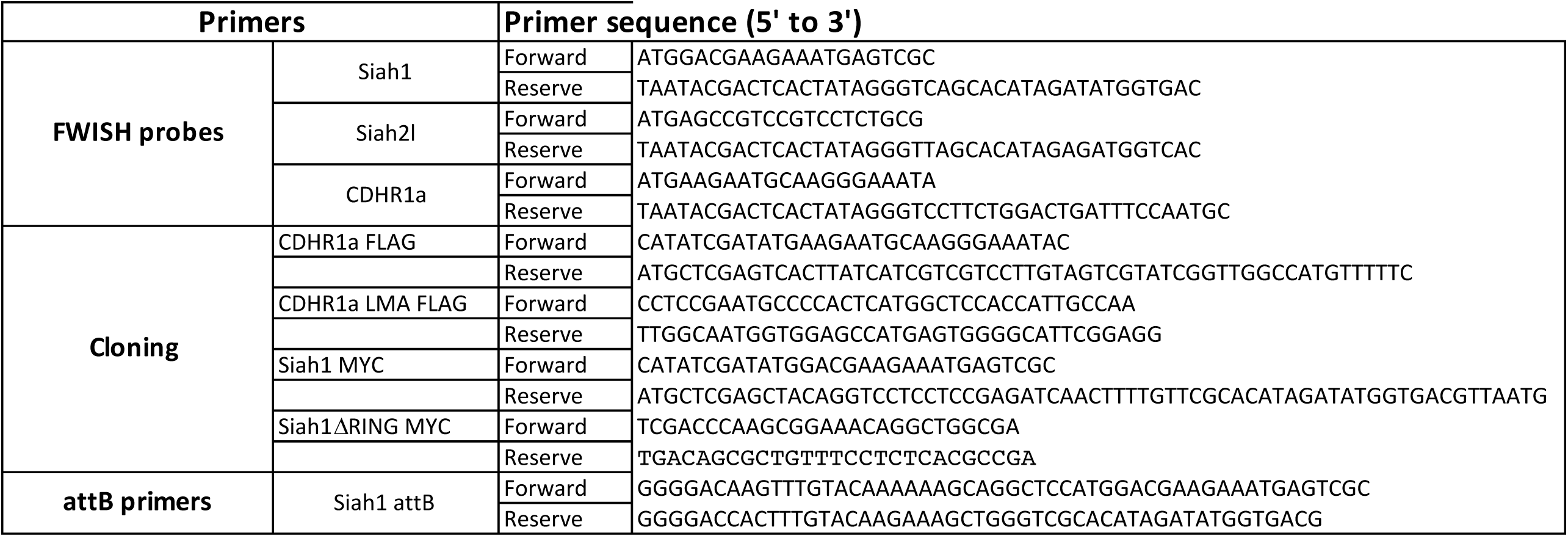

## Material and Methods

### Zebrafish husbandry and embryo maintenance

Zebrafish husbandry used in all procedures were approved by the University of Kentucky Biosafety office as well as IACUC Policies, Procedures, and Guidelines (IACUC protocol 2015-1380). The AB strain was used as wildtype. Transgenic lines used to visualize rod and cone photoreceptors were: Tg[XlRho:EGFP] (XOPS:GFP)^36^ and Tg[3.2TaC:EGFP] (TαC:EGFP)^35^ respectively. Embryos were kept at 28^°^C in E3 embryo media. At indicated times, embryos or larvae were anesthetized in tricaine and fixed with 4% PFA in PBS overnight at 4°C.

### Fluorescent whole-mount in situ hybridization (FWISH)

Fluorescence whole-mount in situ hybridization was performed as a modification from previously described^52^. Embryos were permeabilized with 10mg/mL Proteinase K for 30 minutes for 3 dpf embryos, 60 minutes for 4 dpf embryos, and 75 minutes for 5 dpf larvae. Digoxigenin (DIG) and fluorescein labeled (FITC) probes were synthesized by using DIG and FITC RNA labelling kit (Roche). Primer sequences are listed in Table S1. Anti-Digoxigenin-AP, Fab fragment (ROCHE) and anti-fluorescein-AP, Fab fragment (Roche) antibodies, Fast blue (SIGMA) and Fast-red (SIGMA) were used to detect the hybridization signal.

### Heat shock inducible transgenic zebrafish lines

Tg[hsp70:Siah1], Tg[hsp70:Siah1ΔRING], were generated by amplification the full coding region (Siah 1: Ensembl transcript ID: ENSDART00000026679.8) from 72hpf zebrafish cDNA. The dominant negative Siah1 construct, Siah1ΔRING, was previously described^34^. Both constructs were amplified with primers (Table S1) containing attB for Gateway cloning into pDONR221 using BP Clonase II (Invitrogen). pDONR221 clones were verified by sanger sequencing (eurofins). Using Gateway LR Clonase II, according to the manufacturer’s protocol, (Invitrogen), pDEST constructs pDestTol2Cgred (red heart marker) (gift from Dr. Allison) were combined with pDONR221 plasmids, the 5’ element heat shock promoter plasmid (p5E-hsp70) and the 3’ element (p3E-polyA) plasmid. Positive clones were verified by sanger sequencing (eurofins). Verified constructs were injected (50pg) along with Tol2 mRNA (100pg) and dextran-red into zebrafish zygotes. Transgenic founder embryos were screened at 48-72 hpf for heart marker fluorescence. Founders were outcrossed to wild-type and screened for germline transmission to create the F1 generation.

### Heat shock experiment design

For heat shock at all developmental stages, groups of 50 embryos were incubated at 38°C for 30 minutes using a recirculating water bath. Embryos were then removed from the water bath and placed back in the 28°C incubator in fresh EB media according to the time schedule outlined in Figure 4B.

### Immunohistochemistry and TUNEL Assay

Immunohistochemistry was conducted as previously described^53^ and imaged on a confocal microscope (Leica SP8, Leica). The following antibodies were used: anti-zCDHR1a (CDHR1a, rabbit, 1:100, Bosterbio, Pleasonton, CA), anti-zSiah1 (Siah1, rabbit, 1:100, Bosterbio, Pleasonton, CA), anti-Huc/D (ganglion and amacrine cells, mouse, 1:40,), anti-PKCα (bipolar cells, mouse, 1:100, Santa Cruz Biotechnology, Dallas, TX), anti-Prox1 (horizontal cells, rabbit, 1:1000, Acris, San Diego, CA), anti-PCNA (cells in S-phase, mouse, 1:100, Santa Cruz Biotechnology, Dallas, TX), and activated caspase 3 (apoptotic cells). Alexa fluor conjugated secondary antibodies (Invitrogen, Grand Island, NY) and Cy-conjugated secondary antibodies (Jackson ImmunoResearch, West Grove, PA) were used at 1:200 dilution and DAPI to label nuclei (1:10,000, Sigma, St. Louis, MO). TUNEL assay was conducted with ApopTag Fluorescin Direct In Situ Apoptosis Detection Kit (Millipore, Billerica, MA) on retinal cryosections according to manufacturer’s instructions.

### MG132 treatment

30 embryos were transferred at 52 hpf (3 hours post heat shock) into a 35 mm petri dish containing 5 mL E3 embryo media plus 12.5 mM MG132 (Sigma-Aldrich), or an equal volume of vehicle (DMSO) until 72 hpf. The treatment was refreshed at 61 hpf, immediately after the second heat shock. At 72 hpf, embryos were anesthetized in tricaine and fixed with 4% PFA in PBS overnight at 4°C and washed with PBS with 0.5% Tween-20 (PBS-T) 3 times for 10 minutes.

### DNA constructs, mRNA synthesis and microinjections

All primers used are catalogued in Table S1. Full coding domain sequences for CDHR1a (Ensembl transcript ID: ENSDART00000026679.8) were amplified and cloned into PCS2+. Cdhr1a^LMA^ was generated by site directed mutagenesis of the WT cdhr1a construct and verified by sequencing (eurofins), then cloned into pCS2+. pCS2-CDHR1a and CDHR1a ^LMA^ plasmids were linearized with NotI (NEB). mRNA was synthesized using SP6 mMessage mMachine kit (Ambion) and purified using YM-50 Microcon columns (Amicon, Millipore). mRNA concentration was quantified using spectrophotometry. The mRNA was diluted using nuclease-free water and embryos were injected into the yolk of the embryo at single-cell stage. 100pg of mRNA was used as indicated in the results section.

### Transfection, co-immunoprecipitation and western blotting

For HEK 293 cells transfections, full coding domain sequences for Siah1, Siah1ΔRING, CDHR1a and CDHR1a LMA were amplified and cloned into pCIG2 using In-Fusion HD cloning Plus (Takara). Primers for Siah1 and Siah1DRING included a MYC tag while primers for CDHR1a and CDHR1a^LMA^ included a single FLAG tag. All constructs were verified using sanger sequencing. The HEK 293 cells were cultured at 37°C in DMEM media until they 80% confluency and transfected using TransIT®-LT1 Transfection Reagent (Mirus) at 37°C for 24h. Where indicated, treatment with 10μM of MG132 for the last four hours of transfection was performed. Western blotting and co-IP were performed as previously described^34^.

### Microscopy

For the FWISH, embryos were mounted in 1% low-melting agarose in a glass bottom fluorodish (World Precision Instruments)) prior to imaging using Nikon C2 confocal under the 20X (0.95na) objective. For Immunohistochemistry, the slides were mounted in 40% glycerol (PBS) with coverslips and then imaged using either a Nikon C2 confocal under the 20X (0.95na) and 60X (1.4na) objectives or Leica SP8 confocal under the 20x (0.7na) objective. Images were adjusted for contrast and brightness using Adobe Photoshop.

### Statistical analysis

Two-factor analysis was done by Unpaired Students t-test using GraphPad (https://www.graphpad.com). Data are shown as mean ± St. dev. By conventional criteria, a P value of less than 0.05 was considered significant. ANOVA analysis was performed using Prism8.

## Acknowledgements

We thank Dr. Jessica Blackburn for providing space and reagents for cell culture work. We also thank Dr. Oliver Vocking for assistance with FWISH protocols. WPP was supported by Brazilian National Council for Scientific and Technological Development (CNPq) under grant number 202970/2014-0 and the Morgan Fellowship from the department of Biology, Uniersity of Kentucky. KTT was supported by the Lyman T Johnson scholarship and department of Biology Merit Fellowship, University of Kentucky. This project was supported by the department of Biology, University of Kentucky startup funds awarded to JKF.

